# Recalibration of path integration in hippocampal place cells

**DOI:** 10.1101/319269

**Authors:** Ravikrishnan P. Jayakumar, Manu S. Madhav, Francesco Savelli, Hugh T. Blair, Noah J. Cowan, James J. Knierim

## Abstract

Hippocampal place cells are spatially tuned neurons that serve as elements of a “cognitive map” in the mammalian brain^1^. To detect the animal’s location, place cells are thought to rely upon two interacting mechanisms: sensing the animal’s position relative to familiar landmarks^2,3^ and measuring the distance and direction that the animal has travelled from previously occupied locations^4–7^. The latter mechanism, known as *path integration*, requires a finely tuned gain factor that relates the animal’s self-movement to the updating of position on the internal cognitive map, with external landmarks necessary to correct positional error that eventually accumulates^8,9^. Path-integration-based models of hippocampal place cells and entorhinal grid cells treat the path integration gain as a constant^9–14^, but behavioral evidence in humans suggests that the gain is modifiable^15^. Here we show physiological evidence from hippocampal place cells that the path integration gain is indeed a highly plastic variable that can be altered by persistent conflict between self-motion cues and feedback from external landmarks. In a novel, augmented reality system, visual landmarks were moved in proportion to the animal’s movement on a circular track, creating continuous conflict with path integration. Sustained exposure to this cue conflict resulted in predictable and prolonged recalibration of the path integration gain, as estimated from the place cells after the landmarks were extinguished. We propose that this rapid plasticity keeps the positional update in register with the animal’s movement in the external world over behavioral timescales (mean 50 laps over 35 minutes). These results also demonstrate that visual landmarks not only provide a signal to correct cumulative error in the path integration system, as has been previously shown^4,8,16–19^, but also rapidly fine-tune the integration computation itself.

---

Path integration is an evolutionarily conserved strategy for self-localization that allows an organism to maintain an internal representation of its current location by integrating over time a movement vector representing distance and direction travelled ^4–7^. Place cells and entorhinal grid cells have been implicated as key components of a path integration system in the mammalian brain^20–22^. Thus, we recorded place cells from area CA1 (Extended Data Fig. 1) in 5 rats as they ran laps on a 1.5 m diameter circular track while foraging for liquid reward dispensed at randomized spatial intervals. The track was enclosed within a planetarium-style dome where an array of three visual landmarks was projected onto the interior surface to create an augmented reality environment (Fig. 1a,b). In contemporary virtual reality systems^3, 23–25^, head- or body-fixed rats fictively locomote on a stationary air-cushioned ball or treadmill. Notwithstanding the flexibility of these systems to manipulate the visual experience of the animal, we built the dome apparatus to instead more completely preserve natural self-motion cues, such as vestibular, proprioceptive, and motor efference copy. This system enabled us to test the *a priori* hypothesis that manipulating the animal’s perceived movement speed relative to the landmarks results in a predictable recalibration of the path integration gain.

**Figure 1.**
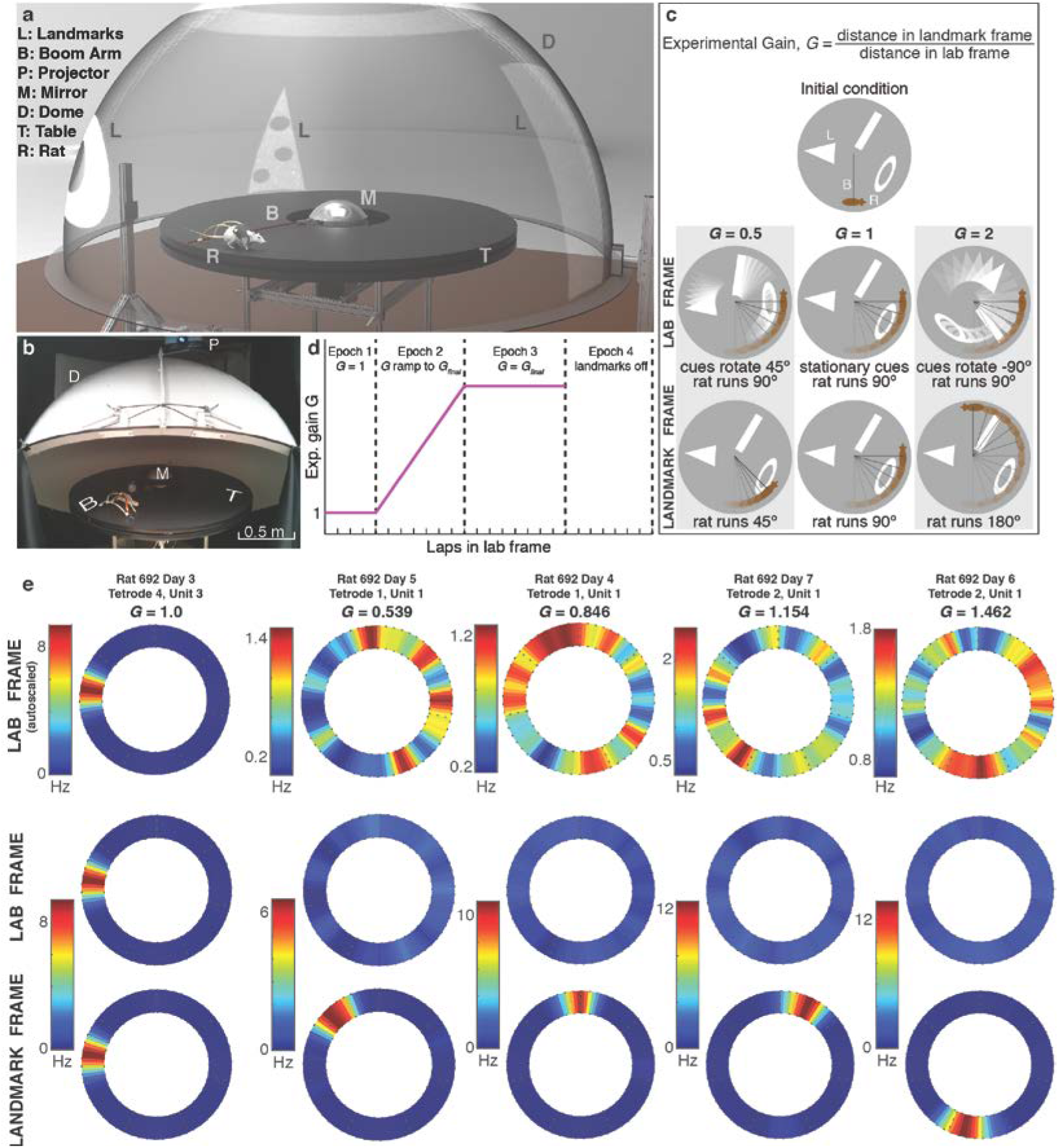
Dome apparatus, experimental procedure, and sample data. **a,** Rendering of dome apparatus. The dome shell is rendered semi-transparent for illustrative purposes. **b,** Photo of the apparatus. The dome is raised in the photo to allow visualization of the interior, but it is lowered as in (a) for the experiment. **c,** Illustration of experimental gain *G*. From the same initial positions of the landmarks and rat, three different gain conditions are shown, in both lab (top) and landmark (bottom) frames of reference. In each case, the rat runs 90° in the lab frame. **d,** Profile of gain change and epochs during a typical session. An annular ring is always projected at the top of the dome (as shown in (a)) for illumination purposes, and is not turned off even in Epoch 4. **e,** Representative firing rate maps for five different units from five separate gain manipulation sessions, shown in the lab frame (top, middle rows) and landmark frame (bottom row) during Epoch 3 (when the experimental gain was constant). The plots in the top row are color scaled to their own individual maximum firing rates, whereas the middle and bottom row plots are color scaled to the maximum firing rate of the bottom plot of each pair. The difference in spatially averaged firing rates between landmark and lab frames results from the distributed firing of the cells over the entire track in the lab frame.

To create the visual illusion that the animal was running faster or slower, the array of landmarks was rotated coherently as a function of the animal’s movement speed. Movement of the landmarks was controlled by an *experimental gain*, *G*, which set the ratio between the rat’s travel distance with respect to the landmarks (landmark reference frame) and its travel distance along the stationary circular track (laboratory reference frame) (Fig. 1c). Recording sessions always began with *G* = 1 (Epoch 1), a control condition where visual landmarks remained stationary alongside the track, so that the rat traveled the same distance in both the landmark and laboratory frames (Fig. 1d). The gain was then gradually changed over the course of multiple laps (Epoch 2) to become less than or greater than one. For *G* < 1, the landmarks moved at a speed proportional to (but slower than) the rat in the same direction; hence, the rat ran a shorter distance in the landmark frame than in the laboratory frame. For *G* > 1, the landmarks moved in the opposite direction; hence, the rat ran a greater distance in the landmark frame than in the laboratory frame. In Epoch 3, *G* was held at a steady-state target value (*G_final_*). In some sessions, the landmarks were then extinguished (Epoch 4) to assess whether the effects of gain adjustment persisted in the absence of the landmarks.

Under gain-adjusted conditions, CA1 units (mean 7.2 ± 5.8 S.D. units/session) tended to fire in normal, spatially specific place fields when the firing was plotted in the reference frame of the visual landmarks, but not when plotted in the reference frame of the lab (Fig. 1e). The strength and continuity of visual cue control over the place fields is highlighted by special cases of *G* (Fig. 2). As *G* was ramped down to 0, the place fields became increasingly large in the laboratory frame (Fig. 2a; Extended Data Video 1). As *G* approached 0, individual units maintained normal place fields only in the landmark frame (Fig. 2b), which resulted in spiking activity that spanned multiple laps in the laboratory frame. At *G* = 0, the animal’s position became locked to the landmark frame, as the landmarks moved in precise register with the rat. Consequently, a unit that was active at that moment would typically remain active throughout Epoch 3, (e.g. yellow unit, Fig. 2a); in contrast, a unit that was inactive at that moment would typically remain silent throughout Epoch 3 (e.g. red unit, Fig 2a). When *G* was clamped at integer ratios such as *3/1* (Fig. 2c) or *1/2* (Fig. 2e), the units maintained the typical pattern of one field/lap in the landmark frame, while firing at the expected periodicity such as 3 times per lap (Fig. 2d) or every other lap (Fig. 2f) in the lab frame. Remapping events sometimes caused different populations of place cells to be active at different times. For example, place cells active during the initial part of the session sometimes went silent (loss of field; Fig. 2e, yellow unit), and place cells silent during the initial part of the session sometimes began firing at a preferred location (gain of field; Fig. 2e, red unit). The remapped cells exhibited normal place fields only in the landmark frame. These examples illustrate that the landmark array exercised robust control over the place fields, outweighing any subtle, local cues on the apparatus as well as nonvisual path integration cues, such as vestibular or proprioceptive cues.

**Figure 2.**
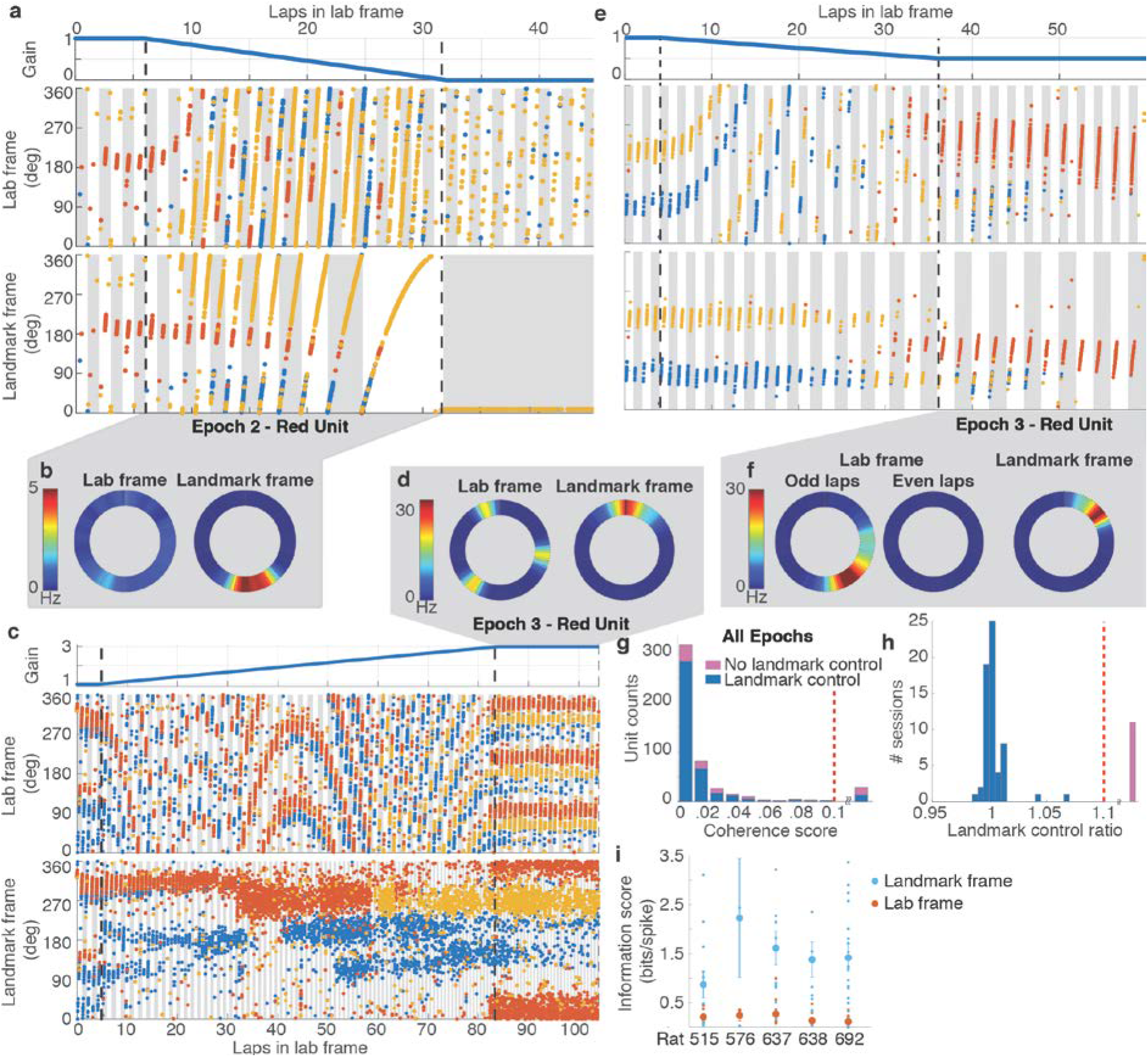
Control of place fields by landmarks. **a,** (top) Profile of experimental gain, *G,* for Epochs 1-3 of a session where *G_final_* was 0. (middle) Colored dots show the location of the rat in the laboratory frame (y axis) as a function of cumulative distance traveled on the track (x axis) when spikes from 3 place cells (red, blue, yellow) were recorded. Alternate gray and white bars indicate laps in this frame. (bottom) The same spikes in the landmark frame. Alternate gray and white bars indicate laps in this frame. The yellow unit fired weakly during the first 8 laps, became stronger on laps 9-10, and maintained the strong field in the landmark frame throughout the remainder of the session. During the last landmark-frame lap, the unit fired as the rat completed ~3 laps in the laboratory frame (middle), thus elongating the place field in that frame to encompass ~1080° of cumulative track angle. **b,** Rate maps of the red unit in laboratory and landmark frames for Epoch 2 of the trial shown in (a). The firing rate is low and diffusely distributed (on average) in the laboratory frame, whereas there is a well-defined place field in the landmark frame. **c,** Epochs 1-3 of a session where the *G_final_* was 2 (same format as (a)). In Epoch 3, all three units maintain normal spatial firing in the landmark reference frame, but they have 3 fields/lap, separated by 120°, in the laboratory frame. **d**, Rate maps of the red unit for Epoch 3 of the trial shown in (c). **e**, Epochs 1-3 of a session where the *G_final_* was 0.5. Remapping occurred near the transition between Epoch 2 and Epoch 3, as the previously silent red unit became active and maintained a stable place field in the landmark frame. In the laboratory frame, however, the unit fired every other lap, (i.e., it was active on the gray laps and silent on the intervening white laps). **f**, Rate maps for the red unit for Epoch 3 of the trial shown in (e). Separate rate maps are shown for the odd- and even-numbered laps in the laboratory frame. **g**, Coherence of the population response. The place fields acted as a coherent population in sessions with (blue) and without (pink) landmark control (see panel h). Units with coherence score above 0.1 (range 0.12 - 0.47) were combined in a single bin (29/500 units). These cells generally displayed poor spatial tuning and therefore did not admit a reliable estimate of hippocampal gain. **h,** Landmark control ratio. In most sessions (blue), the landmark control ratio was ~ 1. Sessions with gain ratio above 1.1 (range 1.16 - 4.02) were combined in a single bin (pink). **i**, Spatial information scores in the lab and landmark frames for each rat. Small dots represent scores from individual units. Mean (large dots) ± s.e.m. are shown.

To quantify the degree of landmark control over the population of recorded place cells, we developed a novel decoding algorithm that was robust to the remapping events described above. We measured each unit’s time-varying spatial frequency (i.e., the frequency of repetition of its spatially periodic firing pattern) from which we then computed a *hippocampal gain*, *H_i_*, for every individual unit, *i*. The median value of *H_i_* over all simultaneously recorded active units during a given set of laps was taken as a population estimate of the hippocampal gain, *H*, for those laps. Just as *G* quantifies the ratio between the rat’s travel distance in the landmark frame versus laboratory frame, *H* quantifies the ratio between the rat’s travel distance in the internal hippocampal “cognitive map” frame^1^ versus the laboratory frame. Hence, if the hippocampal cognitive map is anchored to the landmark frame, then the experimental and hippocampal gains should be identical (*G* = *H*). An ensemble coherence score for each unit was computed as the mean value over the session of | 1 - *H_i_ / H* |, measuring the deviation of *H_i_* from *H* (Methods). The distribution of coherence scores (Fig. 2g) shows that *H_i_* was within 2% of *H* for 80% (399/500) of individual units, and deviations >5% were rare. Even when individual cells remapped, they still exhibited spatial periodicity at gain factors *H_i_* which were close to *H* (see red and yellow units in Fig. 2c). Hence, the population of place cells acted as a rigidly coordinated ensemble from which a precise estimate of *H* could reliably be computed, despite occasional remapping by some place cells.

We quantified the degree of cue control in each session by computing the mean ratio *H/G* for Epochs 1-3 of a session; a ratio close to 1 indicates that the cognitive map was anchored to the landmark frame. The majority of sessions (83.33%, 60/72) exhibited *H/G* near 1, but the rest showed substantially larger *H/G* (> 1.1) indicating loss of landmark control (Fig. 2h; Extended Data Fig. 2). For sessions with *H/G* < 1.1, the spatial information per spike in the landmark frame far exceeded that in the laboratory frame (Fig. 2i; paired t-test, n = 5 rats, t_4_ = 6.213, p = 0.0034). We restricted further quantitative analyses to these sessions, which we defined as demonstrating ‘landmark control’. These results indicate that the augmented reality dome was successful in producing the desired illusion by strongly controlling the spatial firing patterns of the hippocampal cells in the majority of sessions (Extended Data Figs. 3, 4).

Despite strong cue control in the majority of sessions, place fields tended to drift by a small amount against the landmark frame on each successive lap (Fig. 3; Extended Data Fig. 5; also visible in Figs. 2a,c,e and 4a,b) leading to total drifts of up to ~80° over the course of a session. When *G* was gradually decreased, place cells tended to fire earlier in the landmark frame with each successive lap, whereas when *G* was gradually increased, place cells tended to fire later with each successive lap. The accumulated drift over each session was linearly correlated with *G_final_* during Epoch 3 for that session (n = 55 sessions, r_53_ = 0.61, p = 7.2 × 10^-7^; Fig. 3c). The direction of this systematic bias was consistent with a continuous conflict between the dynamic landmark reference frame and a path-integration-based estimate of position (although we cannot rule out the possible contribution of subtle uncontrolled external cues on the track or in the laboratory). That is, when path integration presumably undershot the landmark-defined location systematically (*G* < 1), the place fields shifted slightly backwards in the landmark frame; conversely, when path integration overshot the landmarks (*G* > 1), the place fields shifted forward. The shift may reflect a conflict resolution that is weighted heavily, but not completely, in the direction predicted by the landmark reference frame.

**Figure 3.**
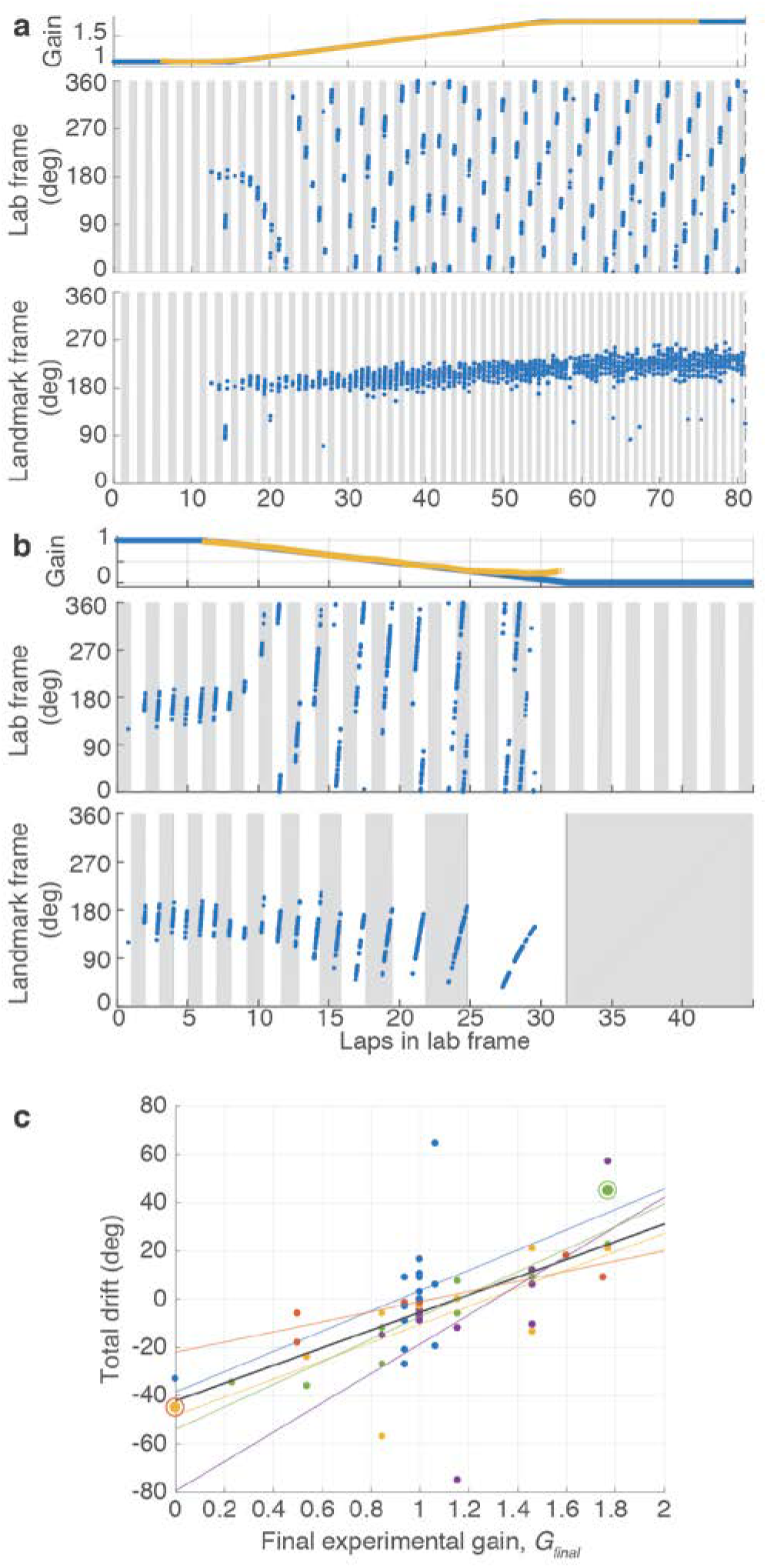
Drift of place fields against landmarks. **a**, Example of positive drift. (top) Experimental gain, *G* (blue) and hippocampal gain, *H* (yellow) for Epochs 1-3 of a session in which *G_final_* was 1.769. There is no *H* (yellow) in the first or last 6 laps due to the 12-lap sliding window. (middle) Spikes from one putative pyramidal cell (blue dots) in the laboratory frame. Figure format is the same as in Figure 2. (bottom) The same spikes in the landmark frame. The unit was silent for the first 12 laps but developed a strong place field in the landmark frame that slowly drifted in the same direction as the animal’s movement over the course of the session. **b**, Example of negative drift from a session in which the *G_final_* was 0. In the landmark frame, the slow drift was in the direction opposite to the animal’s movement direction. Note that the unit was completely silent in Epoch 3, because the rat was not in the place field of the unit as *G* reached 0. **c**, Drift over the entire session vs. *G_final_*. Each point represents an experimental session. Linear fits are shown for each individual rat (colored lines) and for the combined data (black line). The two example sessions of (a) and (b) are shown with the circled markers.

Given the apparent influence of the path-integration circuit revealed by systematic place field drift even in sessions with strong landmark control, we tested whether anchoring of the cognitive map to the gain-altered landmark frame induced a recalibration of the path integrator that persisted in the absence of landmarks. Such recalibration would be evidenced by a predictable change in the hippocampal gain *H* when visual landmarks were extinguished (Fig. 1d, Epoch 4). If the path integrator circuit were unaltered, one would expect that the place fields would revert to the laboratory frame (*H* ≈ 1). Alternatively, if the path integrator gain were recalibrated perfectly, one would expect instead that the place fields would continue to fire as if the landmarks were still present and rotating at the final experimental gain (i.e., *H* ≈ *G_final_*). We found that the hippocampal representation during Epoch 4 was intermediate between these extremes (Fig. 4a,b; Extended Data Video 2): there was a clear, linear relationship between *G_final_* and the hippocampal gain *H* estimated during the first 12 laps after the landmarks were turned off (n = 38 sessions, r_36_ = 0.94, p = 7.9 × 10^-19^, Fig. 4c). Moreover, this linear relationship was maintained when *H* was estimated during the next 12 laps (n = 18 sessions, r_16_ = 0.87, p = 3.37 × 10^-6^, Fig. 4d). The values of *H* for the first and second 12 laps were highly correlated (n = 18 sessions, r_16_ = 0.972, p = 1.72 x 10^-11^, Fig. 4e) with a slope near 1. Thus, *H* was stable over at least 18 laps (i.e., the middle of the second estimation window). Despite this overall stability, there were still fluctuations in *H* in the absence of landmarks (Fig. 4f, Extended Data. Fig. 6). We tested whether changes in behavior could account for the hippocampal gain recalibration by computing several behavioral measures for each epoch, such as running speed and number of pauses/lap (see Extended Data, Behavioral Analysis). Multiple regression analysis showed that *G_final_* strongly predicted *H*, whereas the behavioral variables had negligible influences on *H* (Extended Data Table 1).

**Figure 4.**
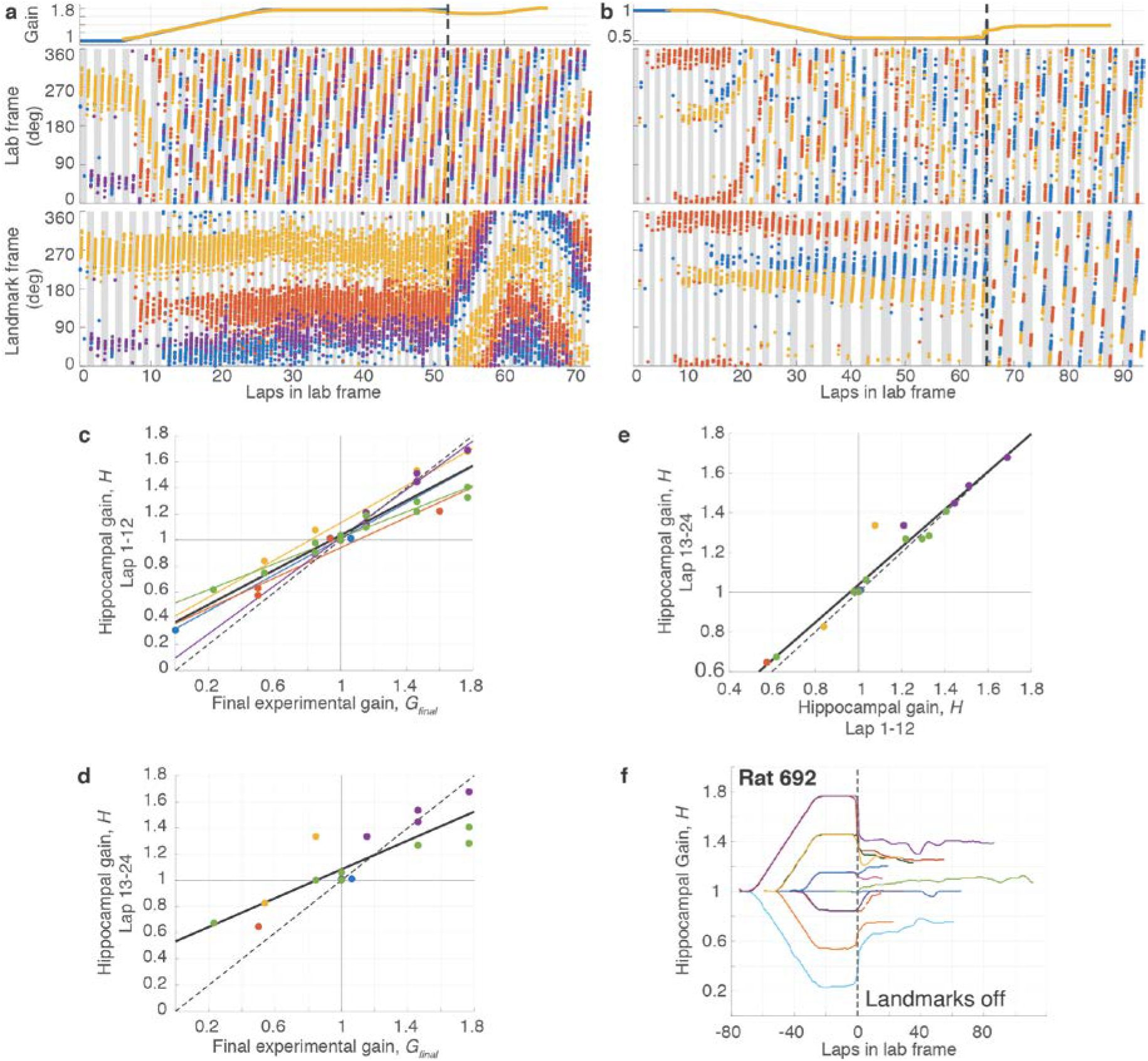
Recalibration of place fields by landmarks. **a**, Example of positive recalibration. (top) Experimental gain, *G* (blue) and hippocampal gain, *H* (yellow) for Epochs 1-3 of a session in which the *G_final_* was 1.769. (middle) Spikes from three putative pyramidal cells (blue, red and yellow dots) in the laboratory frame. (bottom) The same spikes in the landmark frame. When the landmarks were turned off (dashed line, Epoch 4), *H* remained close to *G_final_*, shown by the slower drift of the place fields in the landmark frame compared to the lab frame. (During Epoch 4, the landmark frame was defined assuming the gain was *G_final_* even though landmarks were off.). Note that the traces of *H* (yellow) deviate from *G* (blue) prior to the landmarks turning off; this is an artifact of the sliding window used in the spectrogram and does not affect the conclusions (see Methods, Visualizing *H*). **b**, Example of negative recalibration. The *G_final_* was 0.539. **c**, Recalibration of place fields. The x-axis is *Gfinal* and the y-axis is *H* computed using the first 12 laps (i.e., the value of *H* at lap 6) after the landmarks were turned off. Linear fits for each animal (color) and for the whole data set (black) are shown, along with the perfect recalibration line (dashed line, black) **d**, Sustained recalibration. The x-axis is the same as (c) and the y-axis is *H* computed using laps 13-24 (i.e., the value of *H* at lap 18) after the landmarks were turned off. The number of data points is lower than in (c) because some experiments ended prior to lap 24. **e**, Stability of recalibration. Comparison of *H* during laps 1-12 vs. *H* during laps 13-24. The linear fit is shown in black. **f**, Complete gain dynamics for one animal. For all sessions from one rat, *H* is plotted as a function of laps run in the lab frame. All the sessions are aligned to the instant when the landmarks were turned off (lap 0). The recalibrated *H* was maintained for as many as 50 laps or more.

Using a novel augmented reality dome apparatus, we show here that the path integration system employs a modifiable gain factor that can be recalibrated to a new value that can remain stable for at least several minutes in the absence of salient landmarks. This sustained recalibration can be detected from the spiking activity of hippocampal place cells. Recalibration of this nature has been described extensively in other systems. The cerebellum plays a key role in recalibration of feedforward motor commands during reaching tasks in artificial force fields and during walking on split-belt treadmills^26^. Similarly, the gain of the vestibulo-ocular reflex adapts to changes in the magnitude of retinal slip caused by magnifying glasses, an effect that persists even after the glasses are removed^27^. As with our own results, this recalibration is not perfect in these motor adaptation tasks; i.e., the gain measured after the training trials are biased towards, but not precisely the same as, the experimental gain implemented during the training trials. To our knowledge, such gain recalibration has not been demonstrated physiologically in cognitive phenomena such as spatial representation and path integration (but see ^15^). The lack of complete recalibration may be due to an insufficient number of training laps during Epoch 3, or may reflect inherent limits on the plasticity of the path integrator gain variable.

It is widely accepted that visual landmarks provide a signal to correct error that accumulates during path integration^28^. The results in this paper demonstrate physiological evidence for a role of vision in the path integration computation itself by providing an error signal analogous to retinal slip in the VOR^27^. Specifically, this error signal fine-tunes the gain of the path integrator^15^, minimizing the accumulation of error in the first place. Although it perhaps would not have been surprising to observe gain recalibration over developmental time scales, the rapid recalibration reported in this paper indicates that the path integration gain is constantly and actively fine-tuned even at behavioral time scales. This fine-tuning may be required to (a) maintain accuracy of the path integration signal under different behavioral conditions (e.g., locomotion in the absence of salient landmarks; locomotion on different surfaces that provide varying degrees of slip and cause alterations in the self-motion inputs to the path integrator); (b) synchronize the different types of self-motion signals (e.g., vestibular, optic flow, motor copy, or proprioception) thought to underlie path integration; and (c) coordinate the discrete set of different path integration gains thought to underlie the expansion of grid scales along the dorsal-ventral axis of the medial entorhinal cortex^12,29,30^. The recalibration might be implemented by changes to the head direction^31^ or speed^32,33^ signals that provide input to a path integration circuit. Alternatively, these representations may be unaltered and the gain changes are implemented by changing the synaptic weights between the inputs and putative attractor networks that perform the path integration^9–11, 13^. The augmented reality system described here will allow the investigation of mechanisms underlying the interaction between external sensory input and the internal neural dynamics at the core of the path integration system.

## Acknowledgments

We thank Bill Nash and Bill Quinlan for technical assistance with the construction of the augmented reality apparatus; Marissa Ferreyros, Macauley Breault, Nick Lukish, Jeremy Johnson, and Douglas GoodSmith for technical assistance in running experiments; Balazs Vagvolgyi for software development for validation via camera tracking, Geeta Rao, Vyash Puliyadi, Cheng Wang, Heekyung Lee, Robert Nickl, and Jonathan Bohren for helpful discussions and technical advice; and Adrian Haith for discussions and comments on the manuscript. This research was supported by NIH grants R01 MH079511 (HTB, JJK), R21 NS095075 (NJC, JJK), and R01 NS102537 (NJC, JJK, FS), a Johns Hopkins University Discovery Award (NJC, JJK), a Johns Hopkins Science of Learning Institute Award (JJK, NJC), a Johns Hopkins Kavli Neuroscience Discovery Institute Postdoctoral Distinguished Fellowship (MSM) and a Johns Hopkins Mechanical Engineering Departmental Fellowship (RPJ).

## Author Contributions

J.J.K., N.J.C., and H.T.B. conceived the study. All authors designed the study. J.J.K. and N.J.C. advised on all aspects of the experiments and analysis. F.S. made key contributions to the analysis and interpretation of the data and provided supervision over data acquisition and analysis. R.P.J. and M.S.M. designed and constructed the augmented reality apparatus, performed neurophysiological experiments, and analyzed the data. R.P.J., M.S.M., N.J.C., and J.J.K. wrote the paper and F.S. and H.T.B. provided critical feedback.

## Supplementary Material

### Methods

#### Subjects

Five male Long-Evans rats (supplier Envigo Harlan) were housed individually on a 12:12 hour light-dark cycle. All training and experiments were conducted during the dark portion of the cycle. The rats were 5-8 months old and weighed 300-450 g at the time of surgery. All animal care and housing procedures complied with National Institutes of Health guidelines and followed protocols approved by the Institutional Animal Care and Use Committee at Johns Hopkins University.

#### Dome apparatus

The virtual reality dome apparatus that we designed for this experiment is similar to a planetarium. The hemispherical dome was constructed from fiber glass (Immersive Display UK, Ltd, Essex, UK). The inside surface was uniformly coated with a 50% reflective paint (RAL7040 grey). A hole (15 cm diam.) at the top of the dome allowed light from a video projector (Sony VPL-FH30) with a long-throw lens (Navitar ZM 70-125 mm) to enter. Visual cues were projected onto the inside surface of the dome (Fig. 1). An annular ring of light was projected onto the top, interior surface of the dome; when the spatial landmarks were turned off in Epoch 4, this ring remained on to provide nondirectional illumination.

An annular table (152.4 cm outer diam, 45.7 cm inner diam.) was centered within the dome. The support legs of the dome and the legs of the table were not visible to the rat during the experiment. A commutator (PSR-36, Neuralynx Inc.) was mounted in the center of, but slightly below, the tabletop. The commutator drum was upward, inverted from the typical, ceiling-mounted installation. A hemispherical first-surface mirror (25 cm diam.; JR Cumberland, Inc, Marlow Heights, MD, USA) was mounted to the commutator drum. The image from the projector was reflected off of the mirror and onto the interior surface of the dome. A radial arm (6 mm carbon fiber rod) extending almost to the edge of the table was attached to the central commutator through a smooth bearing. The angle of rotation between the arm and the commutator drum was monitored by a built-in optical encoder. A microcontroller actuated a stepper motor attached to the commutator drum to maintain this angle close to zero, effectively rotating the drum of the commutator along with the radial arm. The rate of rotation of the motor, and correspondingly its auditory noise frequency, was proportional (up to a saturation point) to the speed of the rat in the laboratory frame. The noise could thus potentially serve as an artificial (learned) self-motion cue. If so, the results indicate either that this cue is inconsequential for path integration updating or it is recalibrated along with the natural self-motion cues (i.e., vestibular, motor copy, proprioception, etc.).

Two 3D-printed ‘chariot’ arms for harnessing the rat were attached to the radial arm near the edge of the table. Other lightweight 3D printed components were sometimes attached to the radial boom arm to affix infrared lights, feeding tubes, recording tether supports, etc. The rat wore a body harness (Coulbourn Instruments, Whitehall, PA, USA), onto which Velcro strips and a magnetic attachment pad were sewn. The magnets helped align the harness to paired magnets attached to the chariot arms and the Velcro strip held the rat in that position relative to the arms. During the experiment, the rat pulled along the arm and the components attached to it. Due to the long lever provided by the radial arm and the smooth bearing attachment to the commutator, the load borne by the rat was minimal.

A liquid reward vial and pump and a battery to power the pump and IR lights were mounted to the commutator drum. The commutator drum was connected to a second optical encoder (Hohner Corp., Beamsville, Ontario, Canada) that measured its angular displacement relative to the table. Hence the angle of the rat in the laboratory frame was the sum of the angle measurement from the two encoders (i.e., the angle of the commutator relative to the table and the angle of the radial arm relative to the commutator). A Hall effect sensor (55100-3H-02-D, Litttelfuse Inc., Chicago, IL, USA) mounted to the table, and a corresponding magnet mounted to the commutator drum, were used for post-hoc detection and correction of any spurious jumps in rat angle. To mask auditory cues emanating from outside the dome during the experiments, white noise was played by a speaker placed centrally underneath the table.

A camera was mounted next to the hole at the top of the dome and was hidden from the animal using an annular, concentrically mounted one-way mirror that encircled the hole, occluding the camera from view. The camera provided an overhead view of each experiment, which allowed observation of the experiments and experimenter intervention when necessary (e.g., if the rat broke free from the harness). During the experiments, synchronized video of the rat’s behavior was recorded. To verify our ability to track rat angle, we tracked the location of the boom arm post-hoc using the video recording. We implemented a template-based tracking algorithm using standard subroutines in the freely available OpenCV library (opencv.org, V 3.2.0). Based on the camera resolution (1024 x 768 for the first two animals and 2048 x 2048 for the last three animals), each pixel was calculated to correspond to <1° of the track. The mean absolute error between the video-based tracking and the encoder-based rat angle was small (mean: 3.60° ± 3.86° S.D.) across all 72 sessions.

#### Training

Over 2-3 days, we familiarized the rats to human contact and to wear the body harness. The rats were placed on a controlled feeding schedule to reduce their weights to ~80% of their *ad libitum* weight, whereupon they were trained to run for food reward (either Yoo-hoo^®^ or 50% diluted Ensure^®^) on a training table in a different room from the experimental room. Reward droplets were manually placed at arbitrary locations on the track in the path of the running rat, and the experimenter attempted to lengthen the average interval between rewards to maintain behavior while prolonging satiation. The rats were then transitioned to automatic feeding, where liquid reward was dropped at intervals that varied over time as the rats’ behavior was shaped to maximize forward movement with minimal pauses. The training setup had a similar radial arm and chariot as the main apparatus, but without the surrounding virtual environment. Once the rats were consistently running 30-40 laps without human intervention on the training table, we moved them into the dome and trained them until they ran 30-40 laps in the presence of stationary visual cues. Training usually took 2-3 weeks.

#### Electrode implantation and adjustment

After training, rats were implanted with hyperdrives containing 6 (2 rats) or 12 (3 rats) independently movable tetrodes. Following surgery, 30 mg of tetracycline and 0.15 ml of a 22.7% solution of enrofloxacin antibiotic were administered orally to the animals each day. After at least 4 days of recovery, we began slowly advancing the tetrodes and resumed food restriction and training within 7 days of surgery. Once the tetrodes were close to CA1 they were advanced less than 40 μm per day. Once the tetrodes were judged to be in CA1, as confirmed by sharp wave/ripples in EEG signals and the presence of isolatable units, and the animal was again running at least 30 laps inside the dome, the experimental sessions began.

#### Neural recording

During sessions, the rat was attached to the chariot arms and a unity-gain headstage was attached to its implanted hyperdrive. The neural signals passed through the commutator and were filtered (600-6000 Hz), digitized at 32 kHz, and recorded on a computer running the Cheetah 5 recording software (Neuralynx Inc., Bozeman, MT). Simultaneously, EEG data from each tetrode was filtered (1-475 Hz), digitized at 33 kHz, and stored on the computer. Pulses sent from the experiment-control computer (see below) were time-stamped and recorded as events on the neural recording computer to allow the post-hoc synchronization of the data streams recorded on the two computers.

#### Experimental control

The NI PCIe-6259 data acquisition system (National Instruments Inc., Austin, TX USA) was used to communicate with the dome apparatus. The experiment control was executed by a custom software system coordinated by the software development framework called Robot Operating System^34^ (ROS, Open Source Robotics Foundation, distributed under the BSD-3-Clause License) on a computer running the Linux Operating System (Ubuntu 12.04, 14.04). The custom ROS-based system received information about the rat’s angular position from the two optical encoders and generated the visual scene using standard open-source OpenGL C++ libraries. The visual scene was deformed to match the optics of the projection system and displayed on the projector mounted above the dome. The experimentally measured time lag between movement of the vehicle and movement of the landmark array was 97 ± 24 S.D. ms. The time lag was due to processing time delays as well as to the frame rate of the video projector (17 ms/frame); the jitter was due to occasional frame drops and inconsistencies in update rate due to momentary computational demands (data not shown). We also computed where the landmarks should have been projected if we had instantaneous control. There was no detectable slippage (drift) between the intended location of landmarks and where they were actually projected. The mean absolute error between these values was small for all sessions in which the landmarks were moving (i.e., non-control sessions) (54/72 sessions; mean: 0.59° ± 0.43° S.D.; max: 1.69°).

Rats were rewarded by automatically dropping liquid reward at pseudo-random spatial intervals in the lab frame. These intervals were picked from a uniform distribution with means (typically 40-80°) specified at the beginning of each session. The mean feeding interval was increased gradually during training to delay satiation and maintain running performance, and was generally constant during each experimental session. The experimenter could also dispense reward manually to encourage running behavior when necessary. All the data, including position of the rat, position of the visual stimuli, reward locations, and the overhead video, were saved during the course of the session.

#### Experimental procedure

On each experimental day, baseline data were recorded from the rat for 20 minutes before and after the session while it slept or rested quietly in a towel-lined dish on a pedestal. These sleep data were used post-hoc to confirm recording stability of single units during the trials. During the sessions, the experimenter went into the dome with the rat and always attached the rat to the harness at the same starting location relative to the landmarks (which always were located at the same locations relative to the laboratory frame). After ensuring that the rat was running with a natural gait, the experimenter left the dome. The progress of the session was monitored using the overhead camera, and the experimenter only interfered in cases when the rat partially broke free of the harness, stopped running for long periods, or was running with an unnatural gait.

The session duration varied depending on the running speed of the rat and on how many laps were planned for that session (e.g., ramps to smaller gain values required fewer laps to run the experiment). On days with short sessions, a second session was sometimes run after a short rest duration. The rat was taken out and placed on the pedestal between sessions, to keep the initial conditions consistent. Except on some days where landmarks were kept stationary for the whole duration of the experiment, we took the rat out of the dome only during Epoch 4 (no landmarks inside dome).

#### Experimental gain selection and gain ramp rates

For the first rat (#515), we chose values of *G* close to 1 (1.0625, 0.9375), in addition to one session with a gain of 0. For the second rat (#576) we typically used gains 0.25, 0.5 and 0.75, which resulted in periodic repetitions of place fields in the lab frame. For the remaining three animals, in order to reduce ambiguity of firing patterns in the laboratory and landmark frames of reference, gains were selected in the form of 1 ± *n*/13, *n* = 2,6,10, resulting in gains of 0.231, 0.539, 0.846, 1.154, 1.462, and 1.769. These values ensured that during Epoch 3 the animal’s position relative to the laboratory and landmark frames of reference only aligned once every 13 laps. We used gain ramp rates during Epoch 2 ranging from 1/128 to 1/26 (gain change per lap). The number of laps in Epoch 1 was different for each rat (4 laps for #515 and #576, 6 laps for #637 and #638, and 15 laps for #692). However, the number of laps in Epoch 1 had no apparent relationship to the degree of cue control when the landmarks started to move (proportion of sessions with landmark control failure: #515: 0/15; #576: 1/9; #637: 4/17; #638: 3/14; #692: 3/17).

#### Data analysis

Data from the two experiment computers were synchronized using the paired pulses, and all data were transformed into the same set of timestamps. For each triggered spike waveform, features such as peak, valley, and energy were used to sort spikes using a custom software program (WinClust; J. Knierim). Cluster boundaries were drawn manually on 2-dimensional projections of these features from two different electrodes of a tetrode. We mostly used maximum peak and energy as features of choice; however other features were used when they were required to isolate clusters from one another. Clusters were assigned isolation quality scores ranging from 1 (very well isolated) to 5 (poorly isolated) agnostic to their spatial firing properties. Only clusters rated 1-3 were used for all quantitative analyses in the main text.

To be included in the quantitative analyses, sessions were required to meet the following criteria: (1) sessions with landmark manipulation were completed and the rat was removed in the absence of landmarks, and (2) there were no major behavioral issues / long manual interventions during the session. For the 72/88 sessions meeting these criteria, spikes that occurred when the rat’s movement speed was < 5°/s (~ 5 cm/s) were removed. For each unit, the number of spikes fired when the rat occupied a 5° bin was divided by the time the rat spent in the bin to compute the firing rate. The firing rate was further smoothed with a Gaussian filter of standard deviation 4°. Single units were classified into putative pyramidal cells and putative interneurons by separating them based on firing rate, spike duration, and the autocorrelation function^35^. Only the putative pyramidal cells were used for the main analyses, and the putative interneurons are described in Extended Data Fig. 7.

Spatial information scores were computed by binning and determining firing rates of spikes in both the laboratory and the landmark frames of reference, as described above. If the occupancy-corrected firing rate in bin *i* is *λ_i_*, then information score is computed as:

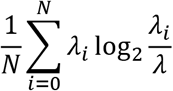

where *N* is the total number of bins, and *λ* is the mean firing rate^36^

#### Behavioral analysis

For each of the 4 epochs, the mean running speed (cm/s), the rate of pauses in running (defined as continuous epochs of 3 seconds or more where the velocity drops below 5 cm/s) (number/lap), the mean duration of each pause(s), the mean interpause temporal interval (s), and the mean interpause spatial interval (cm) were calculated. Interpause intervals were spatial or temporal differences between pause events, where the beginning and end of an epoch were also considered pauses. We first tested whether there were significant changes in these variables between Epochs 1 and 3 (i.e., before and after the gain ramp) and between Epochs 3 and 4 (i.e., before and after the landmarks were extinguished). Next, to address whether changes in behavior predicted the hippocampal gain change in Epoch 4, we ran 2 multiple regression analyses. First, we subtracted the values of each of the behavioral variables in Epoch 1 from the values in Epoch 3. A multiple regression was run with the hippocampal gain (*H*) in Epoch 4 as the dependent variable and the five Epoch 3 – Epoch 1 behavioral measures, as well as the experimental gain (*G*) of Epoch 3, as the regressors. Second, we ran a multiple regression (similar to that above) with Epoch 4 – Epoch 3 behavioral measures, as well as the experimental gain (*G*) of Epoch 3, as the regressors.

#### Estimation of hippocampal gain, *H*

A rat’s position can be decoded from a population of simultaneously recorded place cells using established techniques^37–39^. However, these techniques use an independent dataset to train an estimator and require that the spatial coding be unchanged during the testing phase. In our experiments, there were often remapping events during the gain manipulation epochs, as some units lost their firing fields and other units, which were previously silent, gain place fields on the track. This remapping was typically not all-or-none; rather, different place fields would appear or disappear at different times in the experiments (e.g. Figs. 2c,e, 4a,b). Although the new place fields changed their firing locations coherently with the existing place fields during the experimental manipulations, extensive remapping causes classic population decoding methods to become less accurate or to fail entirely. To solve this problem, we took advantage of the periodicity of firing of the place fields as the rats ran laps on the circular track to measure the *spatial frequency* of the population representation. This spatial frequency is insensitive to the specific place cells that are active at any given moment and it thus forms the core of a spectral decoding technique robust to remapping (Extended Data Fig. 8).

The frequency estimate is termed the ‘hippocampal gain’, *H*. A typical place cell with a single field on a circular track exhibits one field/lap, and hence *H* should be 1/lap (Fig. 1e). As the visual landmarks are moved at an experimental gain *G*, the rat encounters each landmark every *1/G* laps. If the place fields are controlled by landmarks, i.e., they fire every lap at the same location in the landmark reference frame, the value that we estimate for *H* should be similar to the value of *G*. For example, when *G* = 1/2, there should be one field every two laps, and thus *H* = 1/2 (Fig. 2c,d), and for *G* = 3, there should be 3 firing fields per lap, and thus *H* = 3 (Fig. 2e,f).

Hippocampal gain is first estimated independently for all well-isolated units (*H*_i_ for the i^th^ unit) that fire at least 50 spikes per session while the rat is running faster than 5°/s. The spatial spectrogram of the firing rate of each unit was computed at spatial frequencies (i.e., the frequency of repetition of its spatial firing pattern per physical lap) between 0.16/lap and 6/lap, using a sliding window of size 12 laps applied at increments of 5°. The spectrogram was further sharpened using the method of reassignment, which can be used when the input signal contains sparse periodic signal sources^40^. The original spectrogram was also thresholded to the mean + *K* times standard deviation (*K* between 1.1 and 2 based on visual inspection of the raw spectrogram) of its power at each spatial window; this thresholding was then applied to the sharpened spectrogram to improve the signal-to-noise ratio of the spatial frequency content.

The spectrogram can have substantial power in the harmonics of the fundamental frequency, requiring a method to reliably find the fundamental. The gain estimation algorithm identified peaks in the autocorrelation of the spectrogram at each spatial window. Since these peaks typically lie at the fundamental frequency and its harmonics, the fundamental frequency should be both the lowest peak and the difference between peaks. If the median of the difference between peaks was an integer multiple of the lowest peak, the lowest peak was considered the fundamental frequency, and all the power in the reassigned spectrogram further than 0.1 Hz from the fundamental was set to zero (if not, the spectrogram was used as-is). This process was repeated for each spatial window. Finally, the maximum-energy trajectory from the reassigned spectrogram was extracted, and this trajectory formed the time-varying gain estimate for that particular unit. In some cases a particular unit did not produce sufficient spiking activity to generate an estimate for a given window; entries for which there was no estimate were set to NaN in MATLAB for computational convenience. The hippocampal gain estimate for each window for the population (*H)* was calculated as the median *H_i_* from all units under consideration. If there were no active units during a given window (all NaNs) then the value for *H* was set to NaN for that window.

#### Visualizing *H*

For each experimental session, *H* can be plotted as a function of angular displacement of the rat (e.g., Fig. 3a,b, Fig. 4a,b). It is important to note that each estimate is correlated with neighboring estimates due to the 12-lap sliding window. Estimates that are 12 laps apart are calculated from independent data. The estimate at any given angular position is "non-causal" in the sense that it uses neural data from ±6 laps centered around that angular position. This creates the illusion that *H* "anticipates" the extinguishing of landmarks (Fig. 4a,b,f, Extended Data Fig. 6, Extended Data Fig. 7a,b). Inspection of the raw spikes readily verifies that this is an artifact, but this artifact does not affect any of the interpretations in this paper.

#### Coherence score

In a session, if a unit, *i*, is part of a coherent population, its gain should equal the hippocampal gain, namely *H_i_* ≈ *H*. Thus for each 12-lap window we computed a coherence error | 1 - *H_i_ / H* | and defined the coherence score as the mean of this quantity over an entire session.

#### Landmark control ratio

In a session, if the hippocampal gain follows the experimental gain, we expect *H/G =* 1. Thus, *H/G* was computed at each overlapping 12-lap window for Epochs 1-3 and the landmark control ratio was defined as the average of this quantity over a session.

#### Analysis of drift

From each session with landmark control, we identified units which had a single, non-remapped firing field in the landmark frame during Epochs 1 - 3. The average landmark-relative firing rate maps of the unit were calculated separately for the duration of Epoch 1 (start of experiment, *G* = 1) and for the last 12 laps before the landmarks were turned off. The cross-correlation between these two firing rate maps was computed as the rate maps were rotated relative to each other. The landmark-relative angle lag corresponding to maximum correlation was considered to be the drift of the unit. For trials with multiple units with firing fields that did not remap during Epochs 1- 3, we took the mean drift over all units to be the drift for that session. In all, this analysis utilized 136 units from 55 days.

#### Analysis of recalibration

We chose sessions with landmark control and at least 12 laps run after the landmarks were turned off (Epoch 4). The recalibrated gain was selected as the value of *H* six laps after the landmarks were extinguished (lap 6 was the midpoint of the first 12-lap window that includes only data from Epoch 4). To examine the decay rate of recalibration, we chose sessions with landmark control and at least 24 laps run in Epoch 4. We compared the recalibrated gain at lap 6 with the value of *H* at lap 18 (the first point at which the 12-lap spectrogram windows do not overlap).

#### Histology

Once experimental sessions were complete, rats were transcardially perfused with 3.7% formalin. The brain was extracted and stored in 30% sucrose formalin solution until fully submerged, and sectioned coronally at 40 μm intervals. The sections were mounted and stained with 0.1% cresyl violet, and each section was photographed. These images were used to identify tetrode tracks, based on the known tetrode bundle configuration. A depth reconstruction of the tetrode track was carried out for each recording session to identify the specific areas where the units were recorded.

#### Statistics

Parametric tests were used to determine statistical significance. Pearson product-moment correlations were used to test the linear relationship between variables. Paired, 2-sided t tests were used to compare information scores in the laboratory and landmark frames of reference, which assumes normality. Wilcoxon rank-sum tests were used to test differences in behavioral variables. To prevent sampling the same cells across days for this analysis, the experimental session with the greatest number of units was chosen for each rat and for each tetrode.

#### Data availability

The datasets used in this study are available from the corresponding author upon reasonable request.

#### Code availability

Custom code was written for analyzing the datasets used in this study, and generating figures for this manuscript. This codebase is versioned, and uses several third party packages whose license files are included with the respective code. Access to the codebase can be provided by the corresponding author upon reasonable request.

---

**Extended Data Figure 1:**
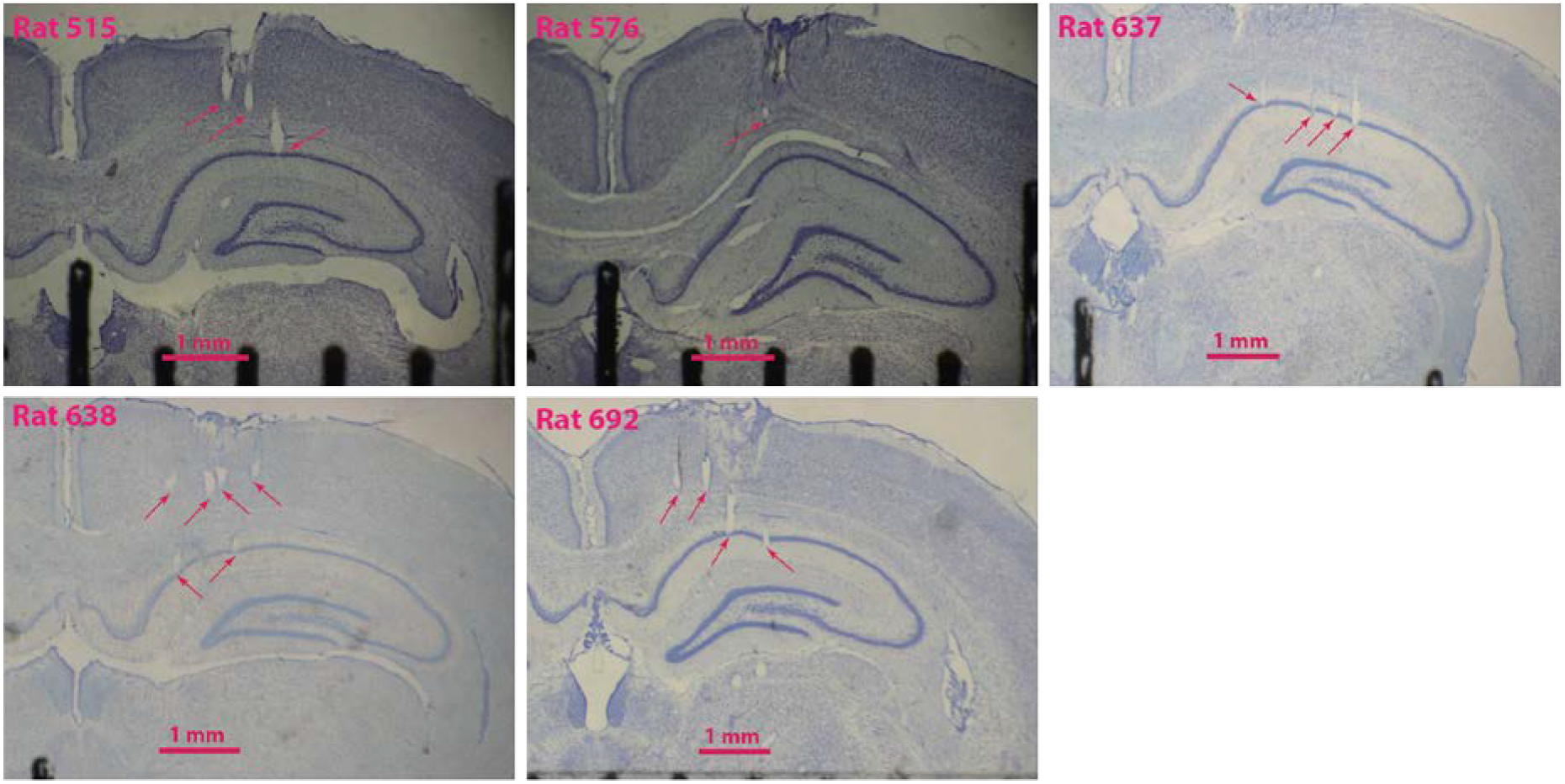
Representative histology. Coronal slices from the five rats used in this study. Arrows point to tetrode tracks in different stages of advancement towards CA1. Note that these are not always the termination of these tetrodes, simply one section along their tracks. In one animal (Rat 576), the histology was inconclusive due to poor fixation and slice quality; however, we determined that the tetrodes were correctly placed in CA1 by the medio-lateral placement of the bundle, tracks in the few sections that we could analyze, and features in the EEG signals observed during recording (e.g., sharp wave/ripples). In one animal, (Rat 638), two of the most medial tetrodes (not shown) appeared to record from the fasciola cinereum, rather than CA1.

**Extended Data Figure 2:**
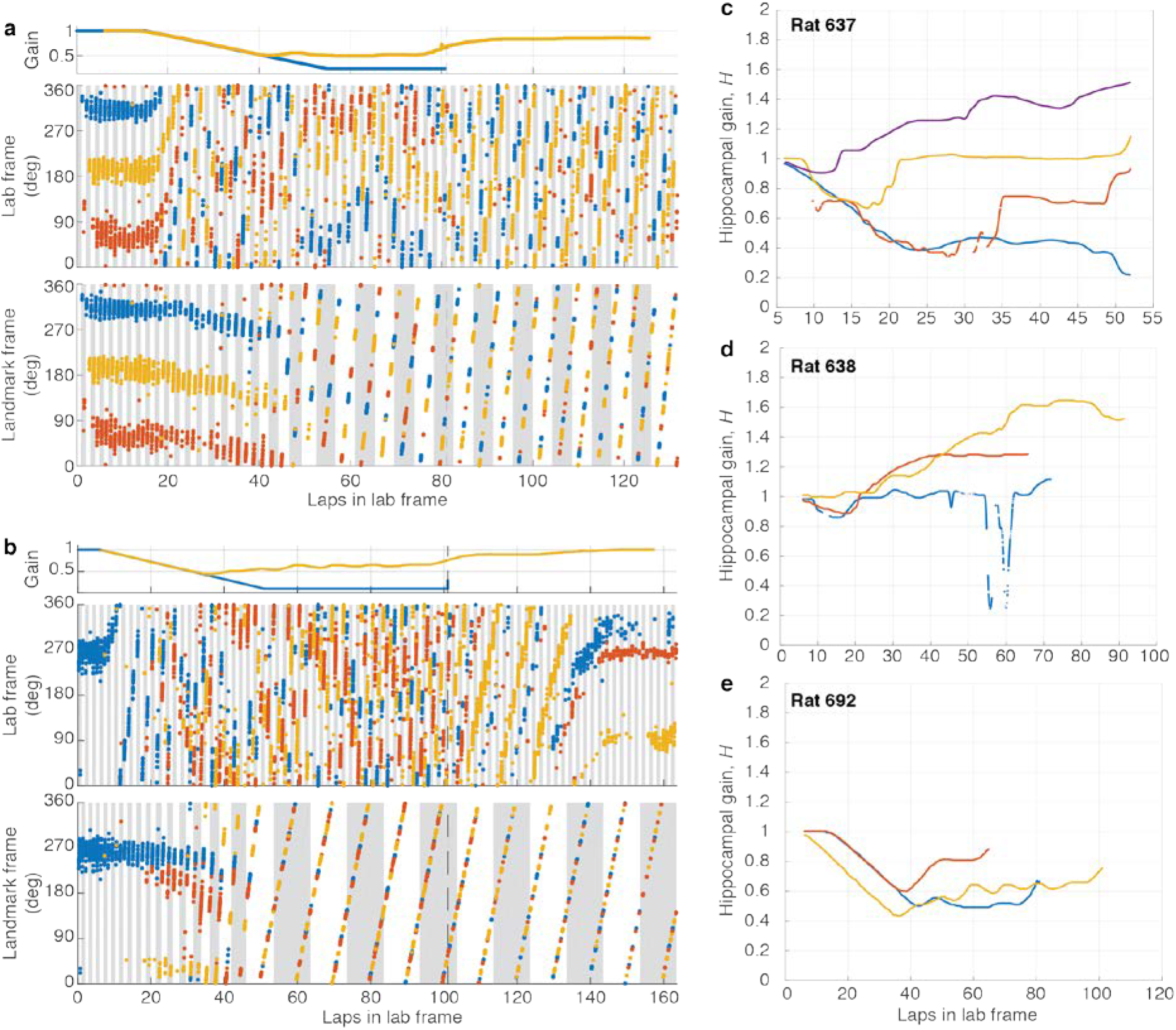
Examples of failure of landmark control. **a**, (top) Experimental gain, *G* (blue), and hippocampal gain, *H* (yellow), for Epochs 1-3 of a session where *G_final_* was 0.231. Note that the two curves overlap until ~lap 40, when they start to diverge. (middle) Spikes from three putative pyramidal cells (colored dots) in the lab frame. Alternate gray and white bars indicate laps in the lab frame. (bottom) The same spikes in the landmark frame. At the point of landmark control failure, the place cells stop firing at a particular location in the landmark frame, and instead start drifting in both lab and landmark frames. Alternating gray and white bars indicate laps in the landmark frame. **b**, Second example, from a different animal, for a session where *G_final_* was 0.1 (same format as (a)). **c-e**, Trajectory of hippocampal gain, *H,* for three rats for all sessions where landmark control failed. The hippocampal gain generally starts near 1 and then diverges from the experimental gain trajectory (not shown) during the session.

**Extended Data Figure 3:**
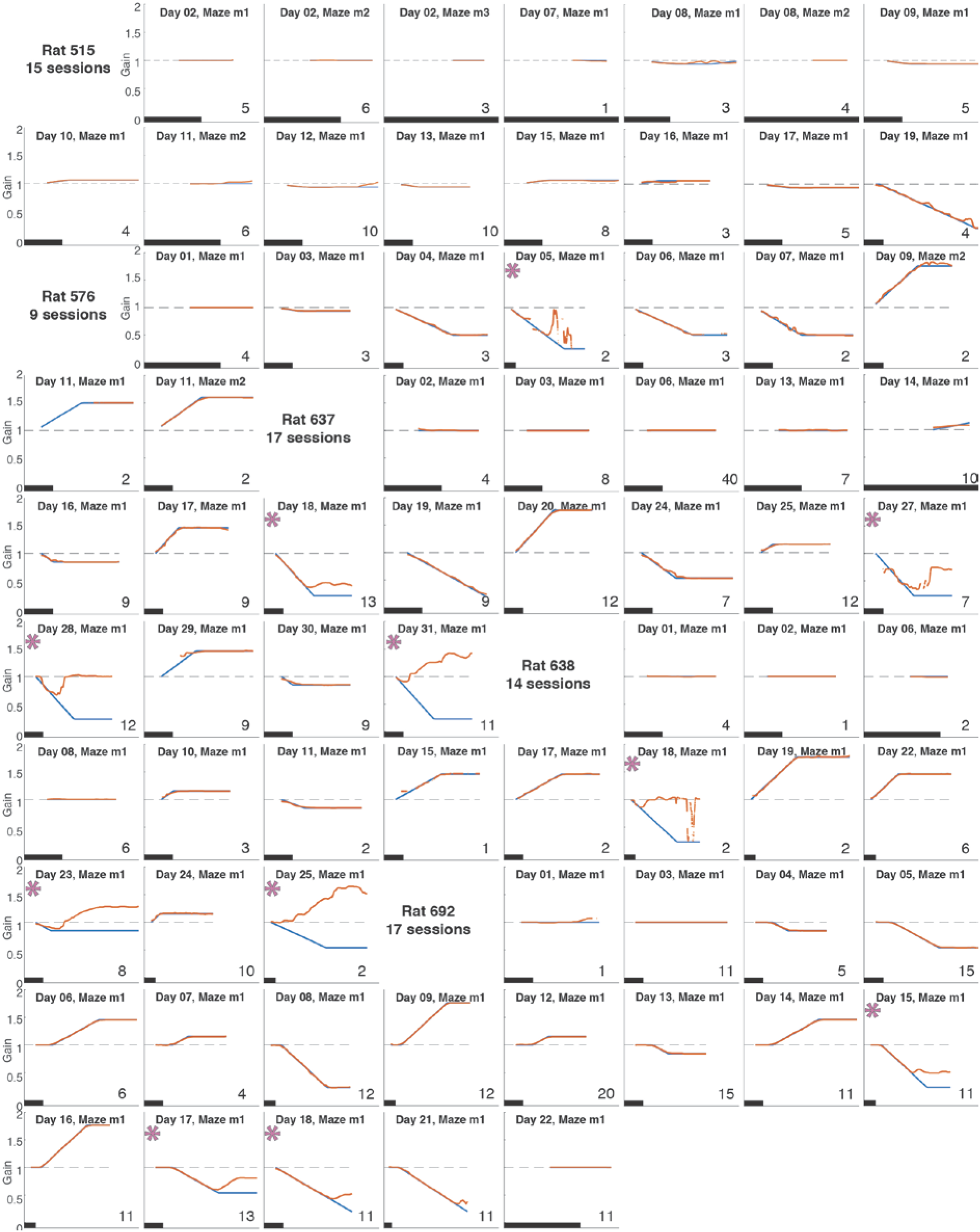
Gain dynamics during each experiment. Each plot represents data from a single experiment. The x-axis is the laps that the rat ran in the laboratory frame (on the table) and the y-axis is gain. The black scale bar in each plot indicates 10 laps. The applied experimental gain (blue) is plotted with the hippocampal gain estimate (red). The ramp rate, length of epochs and final experimental gain for each session can be observed from the curves. An asterisk indicates experiments with loss of landmark control (gain ratio greater than 1.1; see Fig. 2h). In the other plots, the blue and red curves overlap indicating control of landmarks over the place fields. Number of units that passed acceptance criteria (Methods) in each session is indicated in the bottom right hand corner of each plot.

**Extended Data Figure 4:**
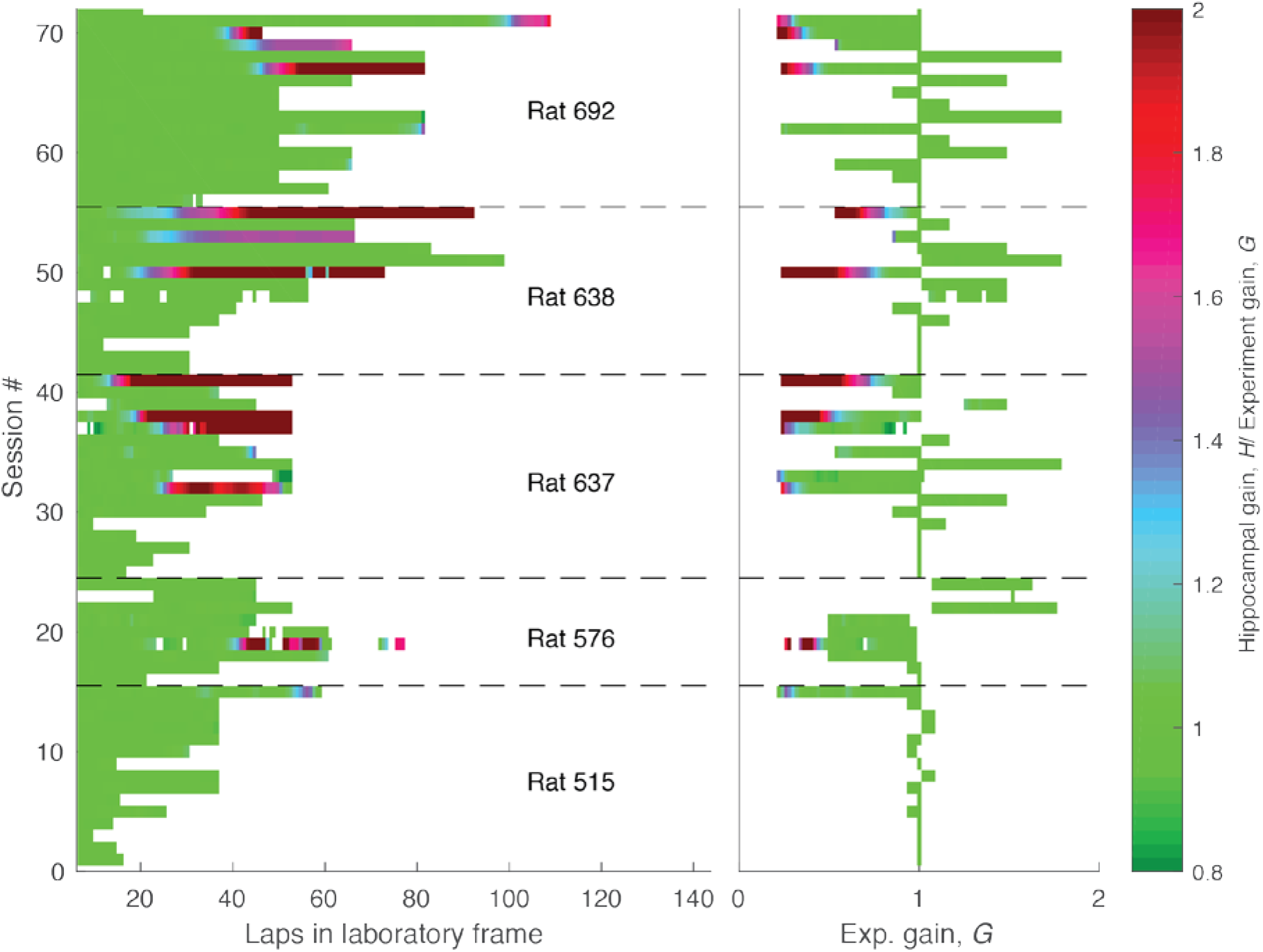
Summary of dataset. Each row indicates one of the 72 sessions composing the dataset during the period when the landmarks were on. In the left plot, the x-axis is laps in the lab frame. In the right plot the x-axis is experimental gain, *G*. The sessions are chronologically ordered (bottom to top). Sessions from different animals are separated by dashed lines. In all rats, we typically performed smaller manipulations in *G* first, since initial landmark failure tended to occur at larger manipulations of *G*. Once landmark control failed, it tended to fail more frequently. The color represents the ratio between hippocampal and experimental gains (*H/G*, color bar, right). Green (*H/G* = 1) indicates landmark control. Four of the rats (576, 637, 638, 692) experienced landmark failure (red portions of trials). Failures only happened when the *G* was less than one (i.e., the landmarks moved in the same direction as the rat) and generally occurred at low values of *G* (less than 0.5) and after rats had experienced multiple gain manipulation sessions over days. The asymmetry in landmark control between *G* < 1 and *G* > 1 is similar to a study of medial entorhinal cortex by Campbell and colleagues^41^. In this study, mice ran on a VR linear track controlled by a stationary treadmill, and the authors manipulated the gain factor between distance travelled on the treadmill versus the VR track. Grid cells showed asymmetric responses to increases versus decreases of the gain. Gain increases (i.e., G > 1) caused phase shifts in the spatial firing patterns but gain decreases (i.e., G < 1) caused changes in the spatial scales. These results were elegantly explained by a model of how grid cells respond to conflicts between self-motion and landmark cues. Although this paper did not address the issues of path integration gain recalibration as in the current study, its results may provide a causal explanation for the asymmetric responses of place cells to the landmark manipulations seen in the present study.

**Extended Data Figure 5:**
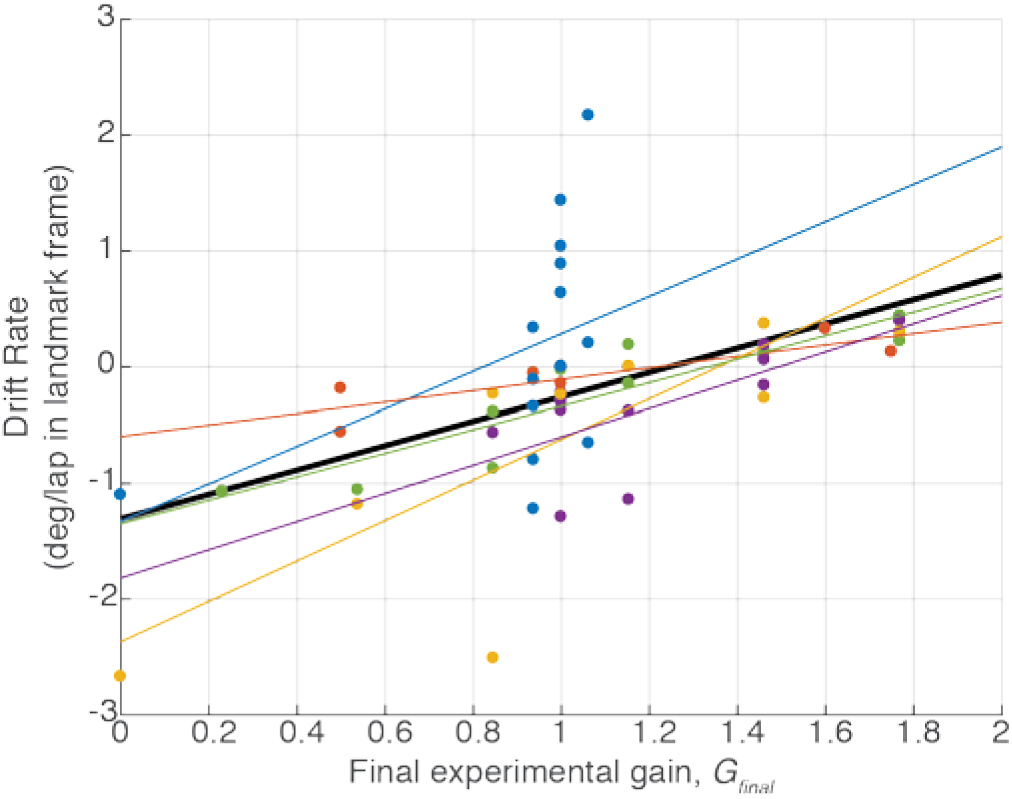
Drift rate vs. Final experimental gain. Figure 3c shows that even in landmark-control sessions, place fields show cumulative drift relative to the landmarks. The magnitude of the drift is correlated with the magnitude of the final experimental gain (*G_final_*). However, a confound is present because the ramp duration in Epoch 2 depends on the value of *G*_final_ (e.g., for *G* > 1, the larger *G_final_* is, the more laps required to ramp *G* up to that value). It is thus possible that the correlation between the total drift and *G_final_* is due to the differences in Epoch 2 duration (and, in some experiments, Epoch 3 duration) rather than due to different rates of drift that depend on *G*. To control for the effect of trial duration, we calculated drift rate by dividing the total drift by the total number of laps in the landmark frame over which the drift was computed. Linear fits are shown for each individual rat (colored lines) and for the combined data (black line; n = 55 sessions, r_53_ = 0.52, p = 4.648 × 10^-05^). These results show that the drift rate was related to the value of *G_final_*.

**Extended Data Figure 6:**
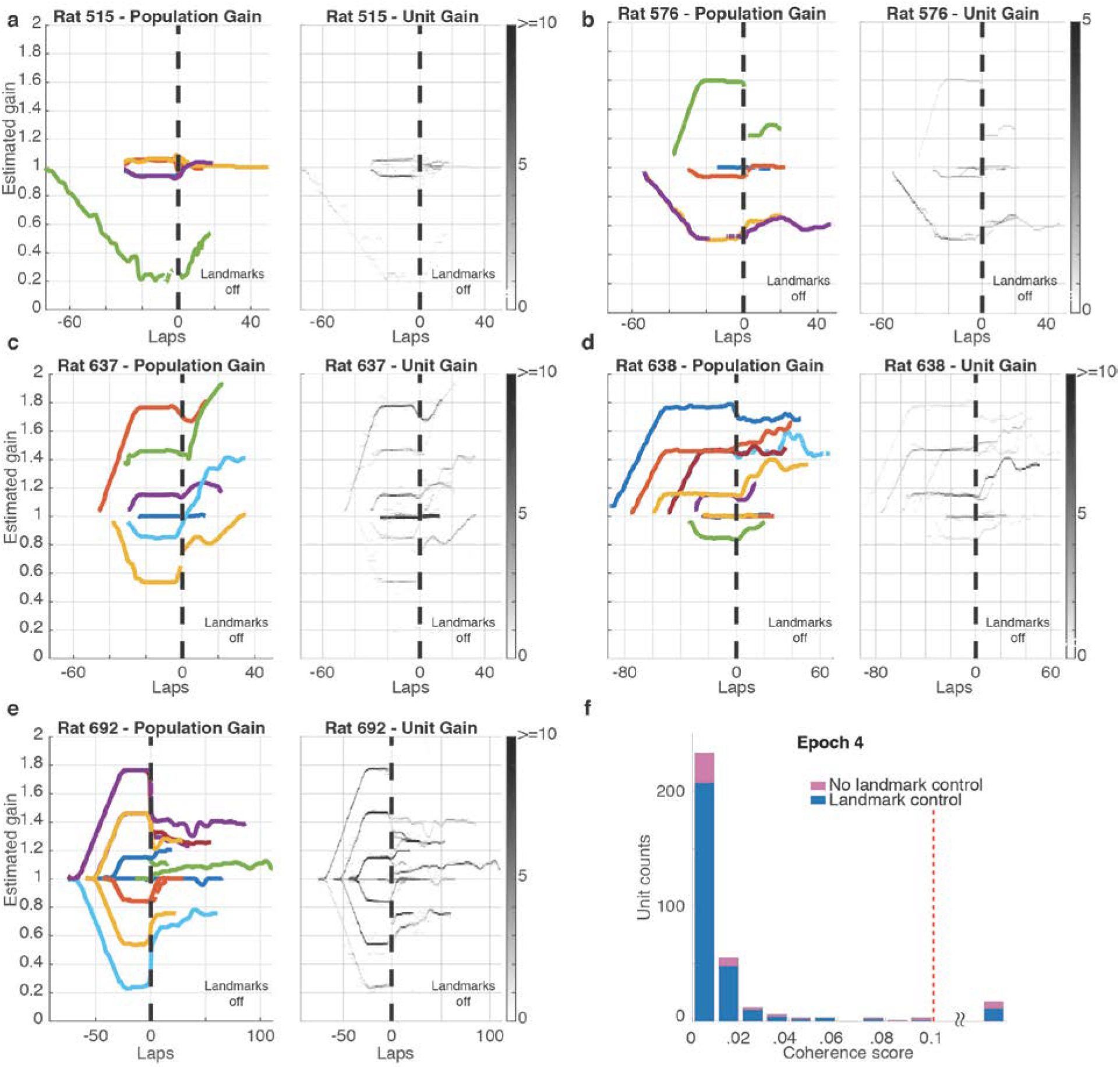
Dynamics of recalibration. **a-e.** The complete hippocampal gain (*H*) dynamics for all 5 rats for trials that exhibited landmark control. (The gain dynamics for Rat 692 is also shown in the main text, Fig. 4e.) In the left panels for each rat (color), *H* is plotted as a function of laps run in the laboratory frame. Sessions are aligned to the instant when the landmarks were turned off (denoted as lap 0). In the presence of landmarks, (before lap 0), the hippocampal gain tracked the experimental gain profiles during a given session (not shown). After the landmarks turned off, the traces largely maintained their recalibrated gain, while also showing some variable drift across experiments. The right panels for each rat show the gain trajectories of all the units in the dataset. The gray scale represents the number of active cells with gains falling in a given bin (bin size is 5° for laps axis and 0.01 for gain axis). These graphs demonstrate the high degree of coherence of the hippocampal population, as almost all cells shared the same gain with minimal deviation. The light-colored lines that occasionally deviate from the main trajectories arise from the small number of cells with poor spatial tuning or from cells that remapped. In the latter case, because our spectral gain analysis used a window of 12 laps, these remapped cells continued to show artefactual values for the limited number of laps that fall in this window but during which the cell was silent. As can be seen, these exceptions had negligible influence on the median population gain values. **f.** Histogram of coherence scores (same format as Fig. 2g) for units firing during Epoch 4 (landmarks off). The shape of the histogram is very similar to Fig. 2g. Almost all units had a coherence score below 0.1, indicating that the place fields acted as a coherent population in sessions with (blue) and without (pink) landmark control in Epochs 1-3, even after landmarks were turned off. Units with coherence score above 0.1 (range 0.11 – 0.41) were combined in a single bin (17/336 units).

**Extended Data Figure 7:**
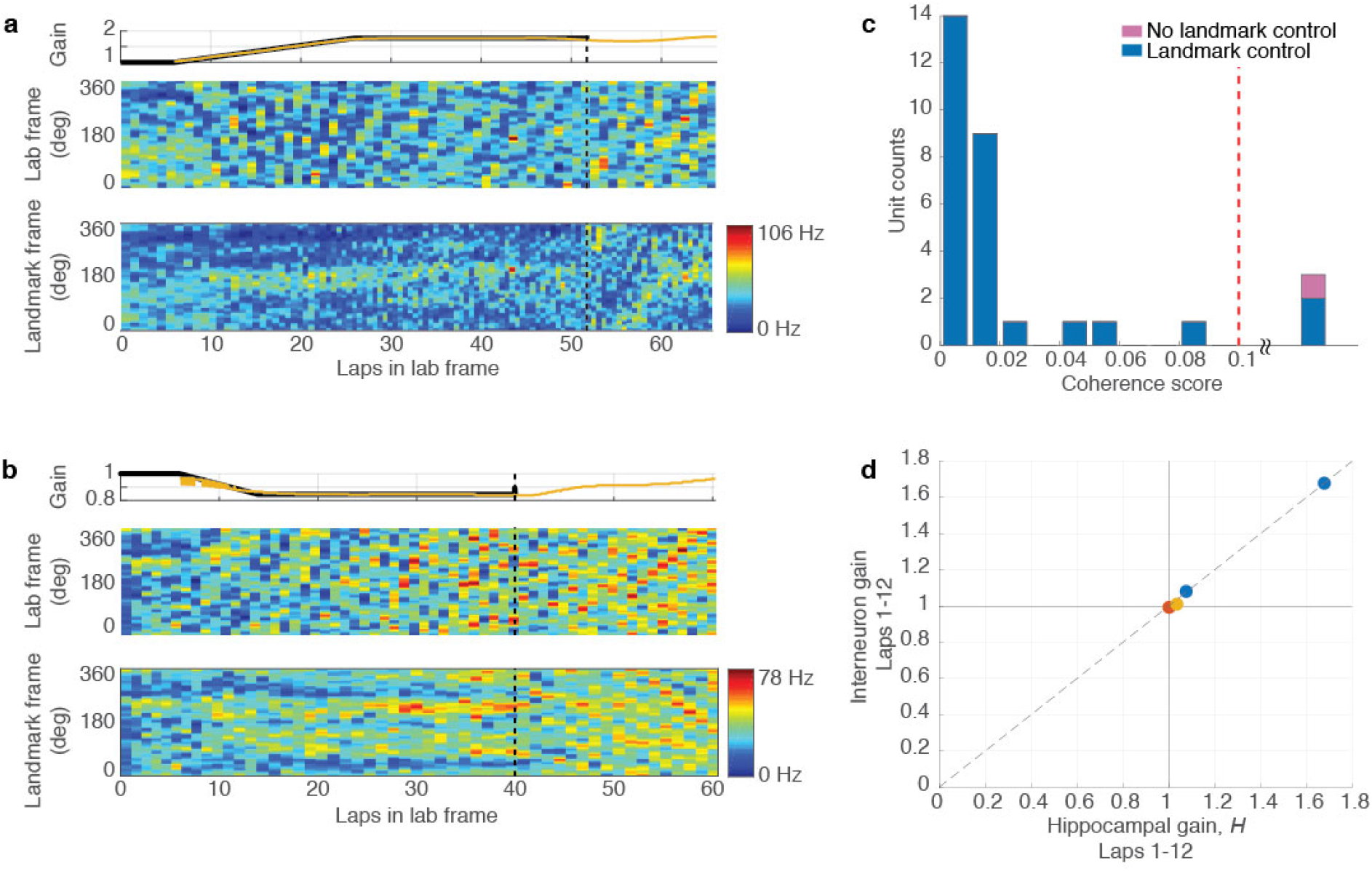
Path integration gain recalibration is also demonstrated by hippocampal interneurons. **a**, (top) Experimental gain, *G* (black) and hippocampal gain, *H* (yellow) for Epochs 1-4 of a session where the *G_final_* was 1.769. *H* was computed as usual from putative pyramidal cells (Methods, Estimation of Hippocampal Gain). In Epoch 4, landmarks are off and hence there is no *G*. (middle) Spatiotemporal rate map of one putative interneuron in the lab frame. Due to the high firing rate of interneurons, rate maps are more illustrative than the spike plots used in place cell examples. Each horizontal bin represents a lap in the laboratory frame, similar to the alternating gray and white vertical bands in the place cell examples (e.g. Fig. 2 a,c,e). Each vertical bin spans 3° in the laboratory frame. (bottom) Rate map of the same unit in the landmark frame. Each horizontal bin represents a lap in landmark frame, and each vertical band spans 3° in landmark frame. Note that the firing pattern is preserved across laps until Epoch 4, when the landmarks turn off. **b**, Example of putative interneuron in a session where *G_final_* was 0.846. Same format as (a). **c**, Histogram of coherence score between interneurons and putative pyramidal cells, as in Fig. 2g. The score for each putative interneuron is computed as the mean value of | 1 – *I / H* | over the entire session, where *I* is the spectral gain estimated from the interneuron, and *H* is the hippocampal gain computed as usual from putative pyramidal cells. Units with coherence score above 0.1 (range 0.15-0.24) were combined in a single bin. **d**, *H* estimated using the first 12 laps after landmarks were turned off, using the median of estimates from putative pyramidal cells compared to the median of estimates from putative interneurons. There are only 4 data points since these are the subset of sessions in Fig. 4c with simultaneously recorded putative interneurons and place cells.

**Extended Data Figure 8:**
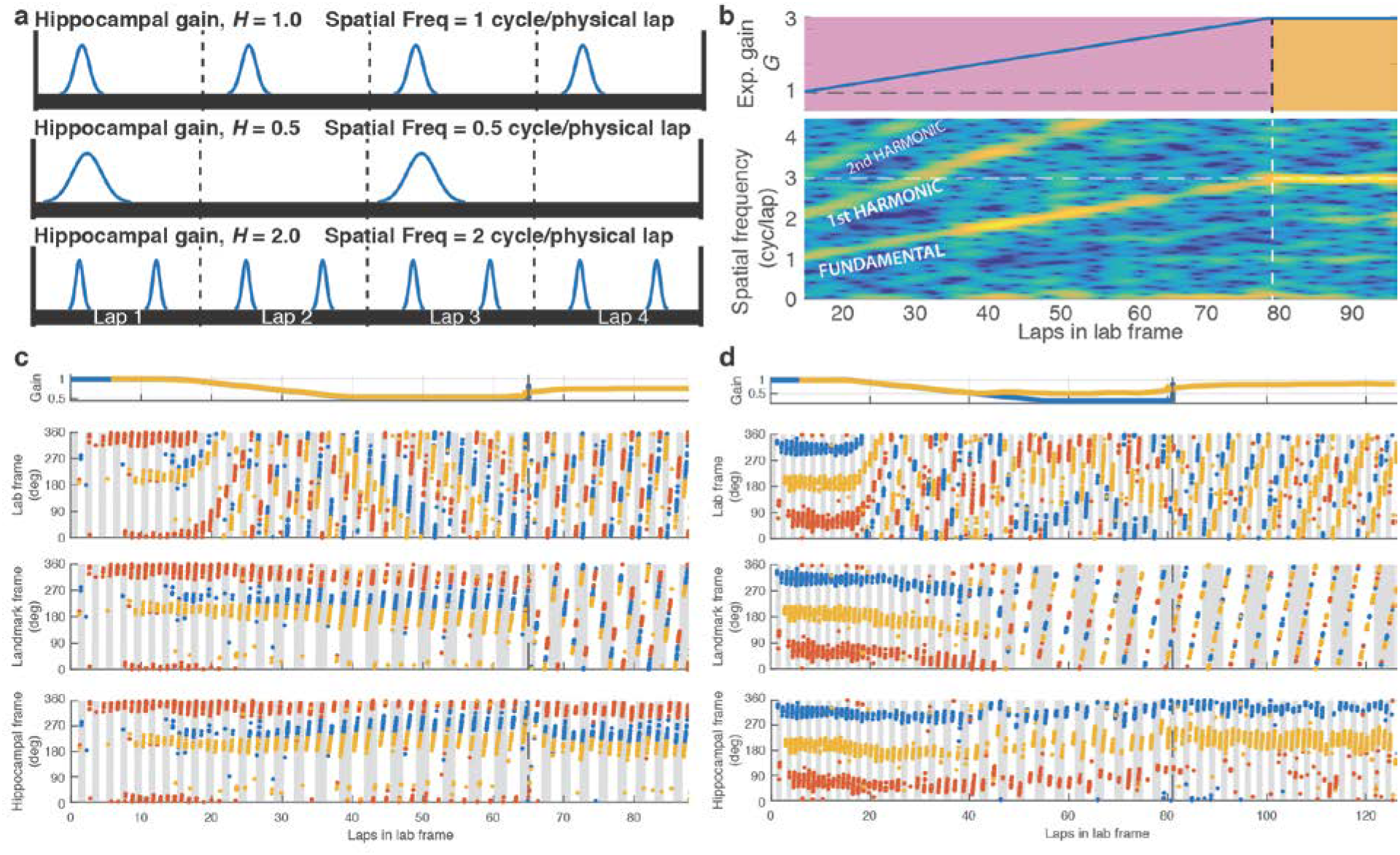
Illustration of spectral decoding scheme. In the dome, as visual landmarks are presented and moved at an experimental gain *G*, the rat encounters a particular landmark every 1/*G* laps (the spatial period). If the place fields fire at the same location in the landmark reference frame, the cell’s firing rate exhibits a spatial frequency of *G* fields/lap. **a**, Illustration of place field firing for three values of hippocampal gain, *H*. **b**, Data from a session in which *G* was gradually increased from 1 to 3 (top) as in Epoch 2 of our experiments. The spectrogram of one unit is shown at the bottom, with the color denoting the power at a given position and frequency. A clear set of peaks in the spectrogram emerges at spatial frequencies corresponding to the experimental gain and at its harmonics. We use a custom algorithm to trace these peaks (Methods, Estimation of Hippocampal gain) and estimate the gain for each unit. The hippocampal gain, *H,* is estimated by taking the median spatial frequency across all isolated units (*H*_i_ for the i^th^ unit) for a given session. Note that this method does not require that cells display single, sharply tuned place fields, as it works for cells with multiple fields as well as for interneurons (Extended Data Fig. 7). **c**, Reproduction of figure 4b, along with addition panel at the bottom that represents the same spikes in the “hippocampal frame;” that is, the spikes were plotted in the frame of the landmarks as if they were rotating at the calculated gain of the place cell map (the hippocampal gain, *H*). The shaded vertical bars denote each lap in the hippocampal frame. Fields from all three units are horizontally aligned in this panel during all epochs, indicating that the spectral decoding technique was successful and that the place fields acted as a coherent spatial representation within the hippocampal frame. **d**, Reproduction of Extended Data Fig. 2a, along with additional hippocampal gain panel at bottom. In this dataset, it can be seen that even after ‘failure’ of landmark control of place fields, the fields are still coherently firing at the same hippocampal gain, which we are able to estimate using spectral decoding.

**Extended Data Table 1:** Results of behavioral analyses.

| | Mean vel<br>(°/sec) | Pauses/lap | Pause<br>Duration<br>(s) | Interpause<br>Interval (s) | Interpause<br>Distance (°) | $G_{final}$ |
| --- | --- | --- | --- | --- | --- | --- |
| *Mean (S.E.M.)* |  |  |  |  |  |  |
| Epoch 1 | 25.1 (0.8) | 0.9 (0.2) | 8.7 (1.0) | 56.2 (9.1) | 924 (156) | -- |
| Epoch 2 | 26.0 (1.0) | 1.0 (0.2) | 6.3 (0.5) | 67.9 (21.3) | 1251 (473) | -- |
| Epoch 3 | 25.8 (1.0) | 1.4 (0.2) | 7.8 (0.5) | 26.7 (4.0) | 483 (91) | -- |
| Epoch 4 | 25.1 (1.1) | 1.5 (0.3) | 9.3 (0.8) | 34.4 (10.9) | 524 (137) | -- |
| Epochs 3-1 | 0.7 (0.5) | *0.5 (0.2) | -0.9 (1.3) | *-29.4 (8.5) | *-441 (134) | -- |
| Epochs 4-3 | -0.7 (0.5) | 0.1 (0.2) | 1.5 (0.9) | 7.7(11.6) | 41 (156) | -- |
| *Multiple regression* |  |  |  |  |  |  |
| <u>Epoch 4 – Epoch 3</u> |  |  |  |  |  |  |
| β | -0.02 | -0.02 | -0.01 | 0.00 | 0.00 | 0.65 |
| S.E. | 0.01 | 0.02 | 0.00 | 0.00 | 0.00 | 0.04 |
| <u>Epoch 3 – Epoch 1</u> |  |  |  |  |  |  |
| β | 0.01 | -0.02 | 0.01 | 0.00 | 0.00 | 0.67 |
| S.E. | 0.01 | 0.03 | 0.00 | 0.00 | 0.00 | 0.05 |

\*Wilcoxon Signed Rank tests were performed on the differences between values in Epochs 3 and 1 and Epochs 4 and 3 with null hypothesis that the difference = 0. Pauses/lap ( $n = 31$ ; $p = 0.0197$ ); Interpause Interval ( $n = 31$ ; $p = 0.001$ ); Interpause Distance ( $n = 31$ ; $p = 0.0035$ ). All other tests for Epochs 3-1 and Epochs 4-3 were not significant

**Extended Data Video 1: Extreme control of place fields by landmarks.** (left) Reproduction of Fig 2a, augmented with moving time marker (vertical dashed line). (right) Overhead videos (8x speed) of the rat running in the dome apparatus as viewed with respect to two distinct frames of reference, synchronized to the time marker in the left plot. Videos show the last ~ 6.5 laps (~ 8 min in real time) of Epoch 2 (*G* ramps to 0). The circular object in the center is the hemispherical mirror (not visible to the rat) used to project images to the inside surface of the dome. Reflections of the three landmarks as well as the annular ring can be seen in the mirror (a small lens artifact also appears on the mirror but was not visible to the animal). Spikes from the same three putative pyramidal cells (red, blue, yellow) are shown at the angular position of the rat. For clarity, spikes only persist for about one lap in their respective frame. (right, top) Original video recorded with respect to the laboratory frame. The yellow place cell is active for ~ 4 laps (over 4 minutes). (right, bottom) Modified video, counter-rotated by the landmark manipulation angle. This results in the reflection of landmarks on the mirror appearing stationary (with a small jitter due to video timestamp resolution). The yellow place cell forms a single field subtending ~180° to 0°.

**Extended Data Video 2: Recalibration of place fields by landmarks.** Same format as Extended Data Video 1. (left) Reproduction of Fig 4a. (right) Videos show approximately the last 4 laps of Epoch 3 (landmarks on) and the first 8 laps of Epoch 4 (landmarks off). (right, top) The place cells do not fire in consistent locations in the laboratory frame. (right, bottom) In the landmark frame, place cells fire in consistent locations during Epoch 3 and then drift slowly in the absence of landmarks during Epoch 4, because the hippocampal gain, *H,* is close to (but not identical to) the final experimental gain, *G_final_*. (During Epoch 4, landmarks are off but the video is counter-rotated as if the gain were *G_final_*.)

